# AURKA promotes malignant properties of neuroblastoma and is downregulated by the PP2A pathway

**DOI:** 10.1101/2025.07.29.665837

**Authors:** Giovanna L. Stadler, Meike Dahlhaus, Frank Speleman, Hans A. Kestler, Miriam Erlacher, Christian Beltinger

**Author notes:** Corresponding author: Eythstraße 24, 89075 Ulm, Germany. The authors declare no potential conflicts of interest.

## Abstract

The role of Aurora kinase A (AURKA) in malignant transformation of sympathoadrenergic progenitors (SAPs), the cells of origin of neuroblastoma (NB), and in progression and maintenance of NB is incompletely understood. *MYCN* is amplified in a subset of high-risk NB cases with poor prognosis. *CDKN2A*, which encodes the tumor suppressors p16INK4A and p14ARF, is deleted or silenced in a subset of NB. PPP2R4 (also known as PTPA) is a key activator of the phosphatase PP2A known to suppress NB.

We show that *AURKA* and *MYCN* expression decreases postnatally in murine adrenal glands, suggesting that their expression may contribute to the maintenance of SAPs. In mouse embryonic fibroblasts (MEFs), used in place of difficult-to-isolate SAPs, AURKA contributed to transformation only when MYCN was overexpressed and INK4A/ARF was depleted. In transgenic mice expressing human AURKA (hAURKA) in adrenal glands, *hAURKA* mRNA was present but protein was undetectable, likely due to AURKA’s short half-life, possibly limiting its role in initial transformation of SAPs. However, AURKA cooperated with MYCN to promote progression of established human SH-EP NB cells lacking INK4A/ARF. In addition, AURKA expression increased during NB progression in TH-MYCN mice and correlated with poor prognosis in human NB, supporting a role for AURKA in disease progression and maintenance. *CDKN2A* and *PPP2R4* mRNA levels also rose during NB progression in TH-MYCN mice, possibly reflecting inadequate tumor-suppressive responses. Bayesian analysis of TH-MYCN expression data supported a role for PPP2R4 in inhibiting NB maintenance. Knockout of *PPP2R4* in KELLY NB cells, along with reduced mePPP2CA, the catalytic subunit of PP2A, increased AURKA protein levels, indicating that PP2A regulates AURKA abundance.

Together, these findings indicate that AURKA promotes malignant properties of NB and is decreased by the PP2A pathway in NB cells

## Introduction

Neuroblastoma (NB), originating from sympathoadrenergic progenitors, is the most prevalent extracranial solid malignancy in children and causes around 15% of all deaths related to pediatric cancers, highlighting its disproportionately high lethality (1). The majority of NB cases are sporadic, frequently exhibiting copy number variations. Among these, *MYCN* amplification represents the most prominent oncogenic driver (1). Overexpression of the transcription factor MYCN transforms immortal fibroblasts (2). Mice with MYCN overexpression targeted to neuroectodermal cells by a tyrosine hydroxylase promoter (TH-MYCN mice) develop NB that are similar to human NB in all major aspects (3–7). Whether MYCN cooperates with AURKA in the initial genesis of NB is unknown.

Aurora kinase A (AURKA) is a serine/threonine kinase essential for mitosis, localizing to the centrosome during interphase and the spindle poles during mitosis. AURKA is tightly regulated and undergoes proteasomal degradation at mitotic exit. AURKA is overexpressed in the majority of high-risk NB (8, 9), where it stabilizes the oncogenic transcription factor MYCN by interfering with SCFbxw7-mediated ubiquitination and proteasomal degradation (10). *AURKA* expression is increased in *MYCN*-amplified NB without being a direct transcriptional target of MYCN (11) and is likewise upregulated in high-risk *MYCN*-non-amplified NB (8). *AURKA* gene amplification is rare in NB (<1%) (8) and no activating mutations have been identified. However, LIN28B and RAN signaling converge on AURKA activity promoting neuroblastomagenesis (12). Taken together, AURKAs role in initial expansion and subsequent malignant transformation of sympathoadrenergic progenitors to NB remains unclear.

Whether altered expression of inhibitors or activators of AURKA contributes to AURKA overexpression in established NB is also not well understood. In particular, the role of Protein Phosphatase 2A (PP2A) has not been defined in this context. PP2A functions as a crucial tumor suppressor across various cancer types and is often found to be downregulated or altered through post-translational modifications (13). PPP2CA is the major catalytic subunit and PPP2R4 (also known as PTPA) is an important activating regulatory subunit of PP2A (14) with a tumor suppressive function (15, 16). Reactivation of PP2A has been investigated as an experimental therapeutic approach to target cancers, including NB (17–19). PP2A decreases phosphorylation of MYCN in NB cells (20) and AURKA protein in Chinese hamster ovary cells (21). Using Boolean network modeling, we previously predicted that PP2A should reduce AURKA protein levels in NB (22).

NB rarely harbor p53 mutations, yet the p14/ARF-MDM2-p53 surveillance axis is disrupted in most cases of NB (23). This dysfunction arises through multiple mechanisms, including BMI1-mediated repression of p14/ARF, reduced activation p14/ARF by decreased CHD5 expression, and MYCN-driven overexpression of MDM2. Additionally, p16/INK4a, which shares the *CDKN2A* locus with p14/ARF, while rarely mutated in NB, as we have demonstrated (24), is frequently inactivated through homozygous deletion, epigenetic silencing and other mechanisms (25–27). Although INK4A/ARF-null mice develop lymphomas and sarcomas (28), they do not develop NB. Whether AURKA-supported transformation and progression of NB requires disruption of INK4A and the ARF-MDM2-p53 pathway concomitant with activation of MYCN, remains unknown.

Here, we show that AURKA cooperates with MYCN to transform *CDKN2A*-deficient MEFs, is upregulated during NB progression in the TH-MYCN mouse model and is decreased by the PP2A pathway in NB cells.

## Materials and Methods

### Isolation and cell culture of wild-type and *CDKN2A*-knockout mouse embryonic fibroblasts (MEFs)

MEFs were generated from day 13.5 post coitum (p.c.) embryos of C57BL/6 mice, 129X/SvJ;B6.129 mice and 129X/SvJ;B6.129-Cdkn2a mice (CDKN2A-knockout) mice following standard dissection and tissue dissociation protocols. Embryonic tissue was minced and incubated with trypsin-EDTA, then cultured in Dulbecco’s Modified Eagle Medium (DMEM) supplemented with 10% fetal bovine serum (FBS), 1% penicillin-streptomycin, and 1% non-essential amino acids (all from GIBCO, ThermoFisher, Waltham, MA, USA). Cells were maintained in 10 cm tissue culture dishes at 37°C in a humidified incubator with 5% CO₂.

### Retroviral constructs and retroviral transduction of MEFs

pWZL-MYCNneo containing the coding sequence of human *MYCN* and pWZLneo as control were kindly supplied by Dr. Martin Eilers, IMT at the Philipps University of Marburg. pBABE-hAurA containig the coding sequence of human *AURKA* was purchased from addgene (plasmid #8510; Watertown, MA, USA) and pBABEpuro as empty control was a gift from Dr. William Weiss, UCSF. Generation of retroviruses and retroviral transduction of wild-type and CDKN2A-knockout MEFs was performed according to standard procedures. Cells were selected with 1.5 µg/ml puromycin (Merck, Darmstadt, Germany) and 600 µg/ml G418 (GIBCO) for at least 10 days.

### Generation of *PPP2R4* knockout in MEFs

sgRNAs targeting exon 1 of the *PPP2R4* gene (NM_002720) was designed using the online platform www.e-crisp.org: sgRNA1 5’-GCAAGATGGCTGAGGGCGAG-3’-CGG (PAM). The sgRNA was cloned into the GeneArt® CRISPR Nuclease Vector (ThermoFisher). MEFs were transiently transfected with the construct, sorted, and seeded as single cells. Single-cell clones were expanded, and genomic DNA was isolated using DirectPCR® Lysis Reagent Cell (Peqlab, VWR, Darmstadt, Germany). Exon 1 of *PPP2R4* was PCR-amplified and subjected to Sanger sequencing. The resulting sequences were analyzed for indels at the target sites. Clones were classified as *PPP2R4* wild-type or knockout. Knockout status was further confirmed at the protein level by Western blot analysis.

### Anchorage-independent growth of MEFs in soft agar

2×10^3^ CDKNA2 knockout MEFs overexpressing MYCN, AURKA, both or neither were seeded in 0.3% top agar that was added onto 0.6% bottom agar in 24-well plates. Low melting point agarose (Invitrogen, Carlsbad, CA, USA) in growth medium containing FBS and additives was used. Growth medium was replaced twice a week until colony formation was observed. Colonies were stained with 1 mg/ml MTT solution (Sigma-Aldrich, Hamburg, Germany) and counted using ImageJ software.

### Subcutaneous transplantation of MEFs

To assess their malignant transformation potential, 2×10⁶ syngeneic MEFs of either wild-type, MYCN-overexpressing, or MYCN and AURKA co-overexpressing phenotype were suspended in 20% Matrigel (BD Biosciences, Heidelberg, Germany) and injected subcutaneously into the right flank of syngeneic C57BL/6 mice. Tumor formation was monitored by palpation twice per week.

### Generation of mice transgenic for *hAURKA* in the sympathoadrenergic system

To generate mice with targeted overexpression of hAURKA in the sympathoadrenergic system, transgenic mice carrying a CAG promoter followed by a floxed chloramphenicol acetyltransferase (CAT) cassette and hAURKA (129X1/SvJ;B6N-Tg(CAG_AURKA(wt))) were crossed with mice expressing Cre recombinase under the control of the dopamine beta-hydroxylase (DBH) promoter (129X1/SvJ-Tg(DBH-iCre)). These are referred to as AURKA mice and DBHiCre mice, respectively. In the absence of Cre, transcription is terminated after the CAT cassette, preventing AURKA expression. In DBHiCre mice, Cre-mediated excision of the CAT-pA cassette enables CAG-driven expression of hAURKA. The DBH promoter drives Cre expression in the peripheral sympathetic nervous system during embryonic and fetal stages, thereby activating AURKA expression in sympathoadrenergic progenitors, the presumed cells of origin of neuroblastoma. Genotyping of recombined alleles was performed using primers flanking the loxP sites, yielding diagnostic PCR products of 2.8 kb for the unrecombined allele and 1.2 kb for the recombined allele. A separate 0.75 kb fragment confirmed the presence of the AURKA transgene. All mouse lines used exhibited no physical or behavioral abnormalities. All animal experiments were performed according to institutional and state guidelines for the protection and care of the animals after approval by the regional authority (Regierungspräsidium Tübingen, Reg.Nr. 986).

### Cell culture of KELLY and SH-EP-MYCN-ER cells

The human NB cell line KELLY was acquired from DMSZ (Braunschweig, Germany) and cultured in RPMI 1640 with 10% fetal bovine serum (FBS), 2 mM glutamine and 100 U/ml penicillin/streptomycin (all from GIBCO) at 37°C in a humidified atmosphere with 5% CO_2_. The cell line has been authenticated by short tandem repeat (STR) profiles using the GenePrint 10 System (Promega, Mannheim, Germany) used according to the manufacturer’s recommendation. SH-EP-MYCN-ER cells have been previously described (29).

### Cooperation of AURKA and MYCN in SH-EP-MYCN-ER cells

SH-EP-MYCN-ER cells were transfected with AURKA expression vector using Lipofectamine 2000 (ThermoFisher) according to the manufacturer’s recommendation. Cells stably selected with 600 µg/ml (GIBCO) were seeded into soft agar in 24-well plates and grown for 2 weeks in medium with 300 nM 4-hydroxytamoxifen (4-OHT, Sigma-Aldrich) exchanged twice per week to translocate MYCN-ER into the nucleus, or with vehicle only. Colonies of more than 30 cells were counted.

### AURKA and CDKN2 in the TH-MYCN NB progression model

The TH-MYCN NB progression model was described previously (30). Briefly, Tg(Th-MYCN)41Waw +/+ mice (3) were sacrificed at postnatal weeks 1 and 2 to isolate sympathetic ganglia containing foci of neuroblast hyperplasia, and at week 6 to collect advanced NB tumors. Sympathetic ganglia from TH-MYCN-/- (wild-type) littermates at weeks 1, 2, and 6 served as controls to monitor gene expression changes during normal sympathetic development. Total RNA was extracted using the RNeasy Mini Kit (Qiagen) following standard procedures. RNA samples were processed and hybridized to SurePrint G3 Gene Expression Microarrays (Agilent, Santa Clara, CA, USA) according to the manufacturer’s protocol. Expression data were summarized and normalized using the vsn method (31) within the R statistical programming environment using the limma package. Linear regression analysis was performed to evaluate the differential temporal expression pattern in ganglia from wild-type mice and ganglia and tumors from transgenic mice. A π-value metric was subsequently calculated as the difference in regression slopes between transgenic and wild-type samples, multiplied with the statistical significance of this difference: π−value = − log10 *p*-value x Δ_slope regression_.

An AURKA expression signature was constructed consisting of expression of AURKA and 18 genes important for regulating AURKA: *AKT1*, *BORA*, *CCDC20*, *CCND3*, *CDH1*, *CENPA*, *CENPE*, *CDK4*, *DLG7*, *KIF11*, *NMYC*, *PLK1*, *PPP1CC*, *PPP2CA*, *PPP2R4*, *PUM2*, *TACC3* and *TPX2*.

### *In silico* analysis of clinically annotated NB patient samples

mRNA expression of *AURKA* and genes of the AURKA network were correlated with clinical outcome by in silico analysis of 498 clinically annotated patient NB (SEQC-GSE62564) using the R2 genomics analysis and visualization platform (https://r2.amc.nl).

### Bayesian analysis of mRNA expression in sympathetic ganglia

Normalized mRNA microarray data from Balamuth et al. (32) were used (GSE17740). 85 genes important in the AURKA network, as deduced from the literature, were subjected to Bayesian analysis. The direction (arrows) and probability (edge thickness) of influence of the top 20 genes according to the differences of mean expression in NB vs. ganglia were shown. Genes were color-coded according to function in relation to AURKA.

### RNA isolation, RT-PCR and qRT-PCR

Total RNA was isolated using TRIzol® reagent (ThermoFisher) or the RNeasy Mini Kit (Quiagen, Hilden, Germany). Purity of mRNA was assessed with a Bioanalyzer device (Agilent). cDNA was synthesized using the SuperScript® III First-Strand synthesis system (ThermoFisher). RT-PCR was carried out with 200 nM dNTPs and 0.1 U/µl Taq DNA Polymerase (both from Sigma-Aldrich). PCR consisted of an initial denaturation at 94°C for 4 min followed by 29 cycles of denaturation at 94°C for 30 sec, annealing at 60°C for 30 sec, and extension at 72°C for 45 sec. The following primers were used: hAurkA forward 5’-ccctgccatcggcacctgaaa-3’, reverse 5’-agctgatgctccactccggctt-3’; hMYCN forward 5’-cgaccacaaggccctcagt-3’, reverse 5’-gggggatgacactcttgagcgga-3’; mAurkA forward 5’-ggcgcgggagagacaaagca-3’, reverse 5’-atgttggggtgccgcaggtg-3’; mMycN forward 5’-gccggaggacaaccccctcc-3’, reverse 5’-gcgccaacctccaactctcccg-3’. Quantitative real-time PCR was performed using the LightCycler Fast Start DNA Master SYBR Green I Kit (Roche, Basel, Switzerland) and a LightCycler 2.0 PCR machine (Roche). Sequences of primers and annealing temperatures were as for RT-PCR. Data was analysed using the LightCycler software version 4.1. (Roche). Relative mRNA expression was calculated by the 2^-ΔCt^ method. The mRNA level of genes investigated was normalized to the mRNA expression of β-Actin.

### Western blot analysis

Cells were lysed in RIPA lysis buffer (Tris (pH8.0, 50mM), NaCl (150 mM), 1% Igepal (NP 40), 0.1% SDS, 1% DOC (sodiumdeoxycholat), EDTA (pH8.0, 1mM), 2 mM DTT (dithiothreitol) and Protease Inhibitor (Roche). Shock-frozen cell pellets were resuspended in RIPA lysis buffer and incubated for 15 min on ice, centrifuged at 16000 *g* and 4°C for 15 min. Protein concentration was measured using the Pierce BCA protein determination kit (ThermoFisher). 20 µg of protein was separated on a Bolt^TM^ 4-12% Bis-Tris Plus gel and blotted onto iBlot^®^ 2 Transfer Stacks (ThermoFisher). Membranes were probed with the primary antibodies rabbit anti-human AURKA (#4718, 1:1000, Cell Signaling Danvers, MA), rabbit anti-human-PPP2R4 (PTPA) (#HPA005695, 1:5000, Sigma-Aldrich), mouse anti-mePPP2CA (sc-81603, 1:500, Santa Cruz Biotechnology, Santa Cruz, CA, USA), mouse anti-Actin (A5441, 1:5000, Sigma-Aldrich) and mouse anti-Tubulin (clone DM1A1, 1:5000, Sigma-Aldrich). As secondary antibodies goat anti-mouse IgG HRP (sc-2005, 1:10000, Santa Cruz Biotechnology) and goat anti-rabbit IgG HRP (G-21234, 1:5000, ThermoFisher, Waltham, MA, USA) were applied. For detection, ECL Western blotting reagent (Amersham, Amersham, UK) and ECLHyperfilms (Amersham) were used. Densitometric quantification of Western blot bands was performed using ImageJ 1.54g (National Institutes of Health, USA).

### AURKA immunohistochemistry

Formalin-fixed and paraffin-embedded slides of mouse adrenal glands and kidneys were deparaffinized and rehydrated. Antigens were retrieved by treating slides for 20 min in 10 mM citrate buffer (pH 6.0) in a pressure cooker. An avidin/biotin blocking system (DAKO, Hamburg, Germany) was deployed, followed by M.O.M. blocking reagent (Vector Laboratories, Newark, CA, USA). Mouse anti-AURKA antibody (#610938, 1:100 in M.O.M dilurent, BD Biosciences, Franklin Lakes, NJ, USA) was applied for 30 min at room temperature, followed by M.O.M. Biotinylated Anti-Mouse IgG Reagent (Vector Laboratories). Samples were treated with streptavidin-alkaline phosphatase (1:500 in TBS, Dianova, Geneva, Switzerland). For detection, the REAL detection system (DAKO) was used. Slides were counterstained with Meyer’s hematoxylin (Sigma-Aldrich) and analyzed using a Keyence BZ-9000E microscope (Neu-Isenburg, Germany) with the Keyence BZ-II Viewer software.

## Results

### AURKA cooperates with MYCN to transform MEFs deficient for INK4a and ARF

Given that NB can arise in peripheral sympathoadrenergic progenitors (SAPs) during postnatal development, we began to investigate a potential cooperation between AURKA and MYCN in neuroblastomagenesis by analyzing their expression in the postnatal mouse adrenal gland. Notably, mRNA levels of both genes declined markedly during normal postnatal adrenal development in mice (Figure 1A).

**Figure 1:**
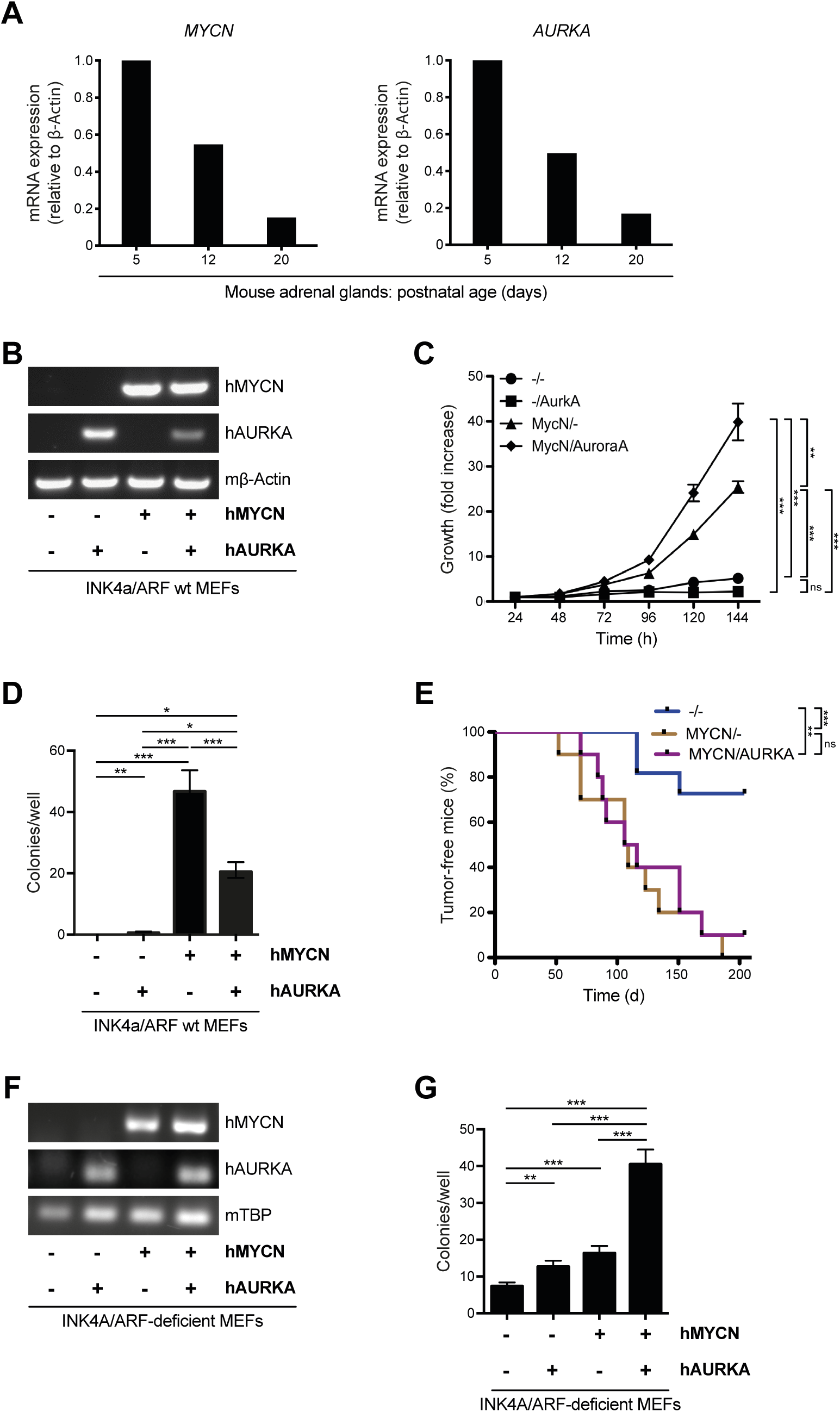
AURKA and MYCN cooperatively transform INK4A/ARF-deficient MEFs. **A) Expression of *MYCN* and *AURKA* mRNA in the murine adrenal gland decreases during postnatal development.** Adrenal glands from three mice were procured at the age indicated and pooled. mRNA was extracted from the pooled glands and qRT-PCR was performed. Expression of *MYCN* and *AURKA* mRNA relative to β-Actin was calculated. **B) hAURKA and hMYCN are expressed in INK4A/ARF wild-type MEFs upon retroviral transduction.** Shown are RT-PCR results. **C) AURKA does not induce proliferation of INK4A/ARF wild-type MEFs while enhancing MYCN-induced proliferation.** Cell proliferation was measured every 24 h by counting cells and is depicted as fold-increase compared to the 24 h cell count. Means of triplicates are shown. ns, not significant; **, *p* < 0.01; *** *p* < 0.001 by Student’s t-test. Similar results were obtained in two independent experiments. **D) AURKA partially inhibits anchorage-independent growth of INK4A/ARF wild-type MEFs overexpressing MYCN.** MEFs were seeded into soft agar in a 24-well plate and grown for four weeks. Colonies of more than 30 cells were counted. Means of triplicates are shown. ns, not significant; *, *p* < 0.05; **, *p* < 0.01; *** *p* < 0.001 by Student’s t-test. Similar results were obtained in three independent experiments. **E) Overexpression of AURKA does not impact on tumorigenicity of INK4A/ARF wild-type MEFs overexpressing MYCN.** Cells in 20% matrigel were injected into the right flank of syngeneic C57BL6 mice (n=8 per group). Mice were palpated for tumor formation twice a week. ns, not significant; **, *p* < 0.01; *** *p* < 0.001 by log rank test. **F) hAURKA and hMYCN are expressed in INK4A/ARF-deficient MEFs upon retroviral transduction.** Shown are RT-PCR results. **G) AURKA cooperates with MYCN to enhance anchorage-independent growth of INK4A/ARF-deficient MEFs.** Stably transduced MEFs were seeded into soft agar in a 24-well plate and grown for four weeks. Colonies of more than 30 cells were counted. Means of sextuplicates are shown. **, *p* < 0.01; *** *p* < 0.001 by Student’s t-test.

We previously demonstrated that the sphere-forming capacity of mouse adrenal tissue, a surrogate marker for the presence of SAPs, decreases after birth (33). Together, these data suggested that sustained expression of MYCN and AURKA may maintain SAPs and promote their malignant transformation. To investigate this notion, we assessed malignant transformation in MEFs, which are readily isolated and experimentally tractable. MEFs overexpressing *hAURKA* alone (Figure 1B) did not proliferate (Figure 1C), failed to grow anchorage-independently in soft agar (Figure 1D), and eventually underwent cell death, precluding their use in syngeneic transplantation assays to test tumorigenicity (Figure 1E). These findings are consistent with previous studies (10, 34). While simultaneous overexpression of *hAURKA* with *hMYCN* (Figure 1B) enhanced MYCN-induced proliferation (Figure 1C), it partially inhibited MYCN-driven anchorage-independent growth (Figure 1D) and had no impact on MYCN-mediated tumorigenicity (Figure 1E). To investigate whether elimination of the G1 checkpoint by disruption of the INK4A and ARF tumor suppressors allows transformation of cells by AURKA alone or in cooperation with MYCN, expression of *hAURKA*, *hMYCN* or both were forced in *Ink4a/ARF* knockout MEFs (Figure 1F). AURKA increased anchorage-independent growth and clearly cooperated with MYCN in this read-out of malignant transformation (Figure 1G).

Taken together, these results indicate that cooperation of AURKA with MYCN in malignant transformation requires primary or secondary dysfunction of the G1 checkpoint, such as disruption of the INK4A and ARF surveillance pathways.

### Mice transgenic for *hAURKA* in the adrenal gland express *hAURKA* mRNA but not protein

To evaluate the potential of AURKA to transform sympathoadrenergic progenitors to NB *in vivo*, *hAURKA* was overexpressed in the sympathoadrenergic system by crossing mice carrying a floxed *hAURKA* allele with mice expressing Cre recombinase under the control of the sympathoadrenergic-specific dopamine beta-hydroxylase (DBH) promoter (Figure 2A). Defloxing of *hAURKA* occurred in the sympathoadrenergic adrenal glands but not in non-sympathoadrenergic organs such as the kidneys of AURKA/DBH-Cre mice, confirming successful targeting of the sympathoadrenergic system (Figure 2B). *hAURKA* mRNA was specifically expressed in the adrenal glands but not the kidneys of AURKA/DBH-Cre mice, indicating successful organ-specific transcription of *hAURKA* (Figure 2C). Surprisingly, hAURKA protein was not detected in the adrenal medulla, the sympathoadrenergic compartment of the adrenal gland (Figure 2D). This absence was not attributable to technical limitations of immunohistochemistry, as hAURKA protein was readily and specifically detectable in human NB samples (Figure 2D). This implies that AURKA is either not translated, or, more likely, rapidly degraded in normal sympathoadrenergic tissues.

**Figure 2:**
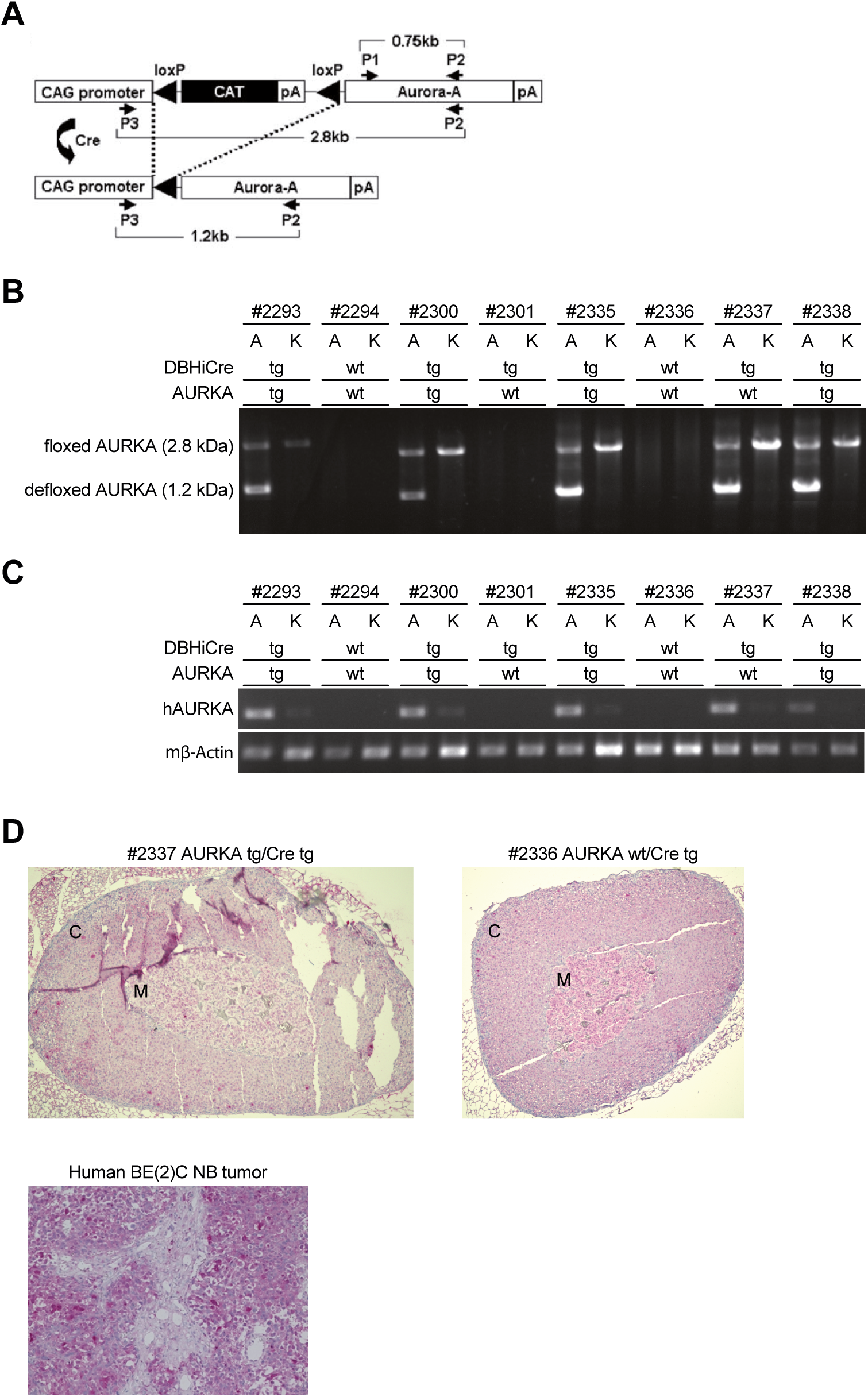
Mice transgenic for *hAURKA* in the adrenal gland express *hAURKA* mRNA but not protein. **A) Schematic of the *hAURKA* transgene and its Cre-mediated recombination in the sympathoadrenergic system.** Shown are constructs, Cre-mediated recombination and genomic analysis of mice and their offspring with floxed hAURKA and Cre recombinase under control of the sympathoadrenergic-specific DBH promoter. CAG, synthetic composite promoter; Cre, Cre recombinase; loxP, recognition site for Cre recombinase; pA, polyadenylation signal; P1, P2 and P3, PCR-primer binding sites. **B) *hAURKA* is successfully defloxed in the adrenal glands of AURKA/DBH-Cre mice.** DNA was isolated from adrenal glands (“A”) and kidneys (“K”) of representative offspring (“#”, ID of offspring) of crossings between mice with floxed hAURKA and mice with DBH-Cre recombinase. Shown are the PCR genotyping results for floxed and defloxed AURKA using primers P2 and P3. “tg” denotes transgenic, “wt” wild-type, “DBHi Cre” Cre recombinase under the control of the DBH promoter and AURKA. **C) *hAURKA* mRNA is specifically expressed in the adrenal glands of AURKA/DBH-Cre mice.** Depicted are the qRT-PCR results for *hAURKA* mRNA using primers P1 and P2, and for mouse β-Actin mRNA as control. hAURKA, human AURKA mRNA; mβ-Actin, mouse β-Actin mRNA. **D) hAURKA protein is not present in the adrenals of AURKA/DBH-Cre mice while readily detectable in human BE**(**2**)**C NB cells.** Shown are representative stainings of hAURKA in an adrenal gland of an AURKA transgenic/DBH-Cre transgenic mouse (AURKA tg/Cre tg; upper left panel), an AURKA wild-type/DBH-Cre transgenic mouse (AURKA wt/Cre tg; upper right panel) and in a human BE(2)C NB tumor growing subcutaneously in RAG-/- cgc/-/- immunodeficient mice (lower panel). C, adrenal cortex; M, adrenal medulla.

### Expression of *AURKA*, *CDKN2A* and AURKA regulators increases during development of NB in TH-MYCN mice

*In vivo* investigations into the genesis of NB are restricted by the inaccessibility of human peripheral sympathetic nervous tissue from which NB originates. In contrast, this tissue and its corresponding NB are readily accessible in transgenic mice. Thus, we probed the AURKA network in NB by reanalyzing *in silico* expression data from the TH-MYCN NB model. Of note, *AURKA* transcription was already upregulated in pretumorous ganglia and increased further during progression to NB (Figure 3A). This suggests that AURKA may be a driver in the development of NB. Concomitantly with expression of *AURKA*, expression of the CDKN2A genes I*NK4A* and *ARF* increased early, which may represent a futile control mechanism by these tumor suppressors during development of NB in this model (Figure 3B). Along this line, expression of *PPP2R4*, an activator of the phosphatase PP2A implied in posttranslational control of AURKA, increased (Figure 3C). Furthermore, an AURKA signature encompassing genes important in AURKA regulation increased early during tumor development (Figure 3D). Taken together, these data support a role of AURKA in murine neuroblastomagenesis.

**Figure 3:**
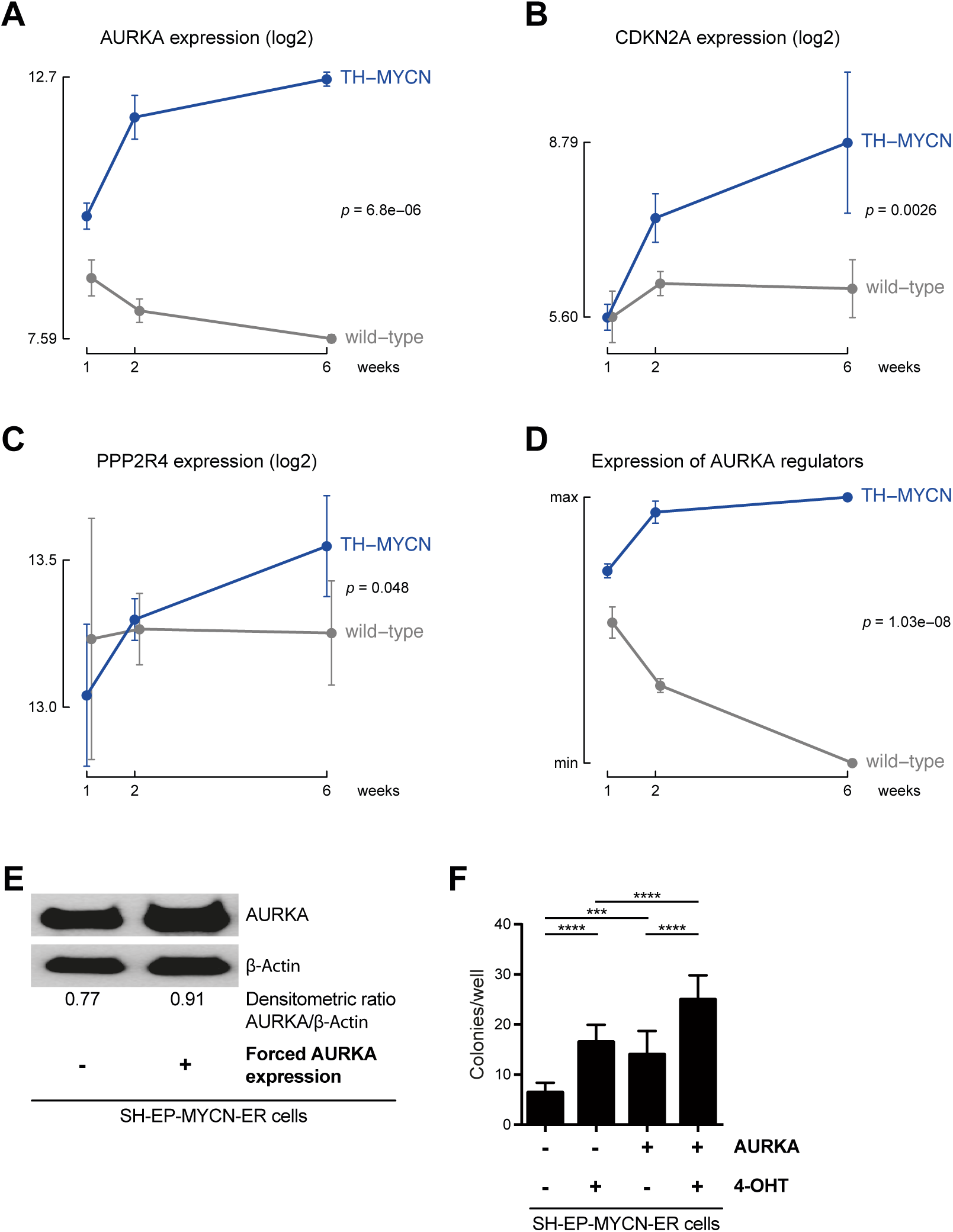
AURKA and its regulators are upregulated during NB development in TH-MYCN mice, and AURKA cooperates with MYCN to enhance malignancy of SH-EP NB cells. Shown is mRNA expression of *AURKA* (A), *CDKN2A* (B), *PPP2R4* (C) and of an AURKA signature **(D)** consisting of expression of AURKA and 18 genes important for regulating AURKA during progression of NB in ganglia of TH-MYCN mice compared to ganglia from wild-type mice. The means and standard deviations of four samples depending on age are depicted. The difference in the regression slope between transgenic and wild-type samples was calculated. The multiple testing-corrected *p*-value for the linear regression analysis is shown. **E) Enhanced expression of AURKA after transfection and stable selection of SH-EP-MYCN-ER cells.** Representative Western blot of AURKA in transfected SH-EP-MYCN-ER cells. AURKA expression was quantified by densitometry relative to β-Actin. **F) Enhanced anchorage-independent growth of SH-EP-MYCN-ER cells overexpressing AURKA upon induced nuclear translocation of MYCN-ER.** Stably selected SH-EP-MYCN-ER cells were seeded into soft agar in 24-well plates and grown for 2 weeks. 4-hydroxytamoxifen (4-OHT) was added to culture media to translocate MYCN-ER into the nucleus. Colonies of more than 30 cells were counted. Means of more than 20 wells are shown. ***, *p* < 0.001 and **** *p* < 0.0001 by one-way ANOVA.

### AURKA cooperates with MYCN to promote anchorage-independent growth of established NB cells lacking INK4A/ARF

Having shown that AURKA cooperates with MYCN in the initial transformation of INK4A/ARF-deficient MEFs and plays a role in murine neuroblatomagenesis, we next investigated whether AURKA enhances malignant properties of established human NB cells lacking *INK4A/ARF*. We used SH-EP-MYCN-ER cells, which carry a homozygous deletion of *CDKN2A* (35) and express MYCN fused to a modified estrogen receptor (MYCN-ER). Upon treatment with 4-hydroxytamoxifen, protein expression of AURKA was moderately increased (Figure 3E). Nuclear translocation of cytoplasmically sequestered MYCN-ER upon addition of 4-hydroxytamoxifen significantly increased anchorage-independent growth (Figure 3F). Moderate overexpression of AURKA alone also enhanced anchorage-independent growth. Notably, combined AURKA overexpression and MYCN activation increased anchorage-independent growth beyond the effect of either alone (Figure 3F).

### High mRNA expression of *AURKA and PPP2R4* is significantly associated with poor prognosis in patient NB

To investigate the role of AURKA and PPP2R4 in the patient setting, clinically annotated data from human NB were analyzed in silico. High mRNA expression of *AURKA* and *PPP2R4* were significantly associated with poor prognosis in patient NB (Figure 4A, left and middle panel). High mRNA expression of *PPP2CA* was also associated with poor survival, although without reaching statistical significance (Figure 4A, right panel). High combined mRNA expression of *AURKA* and *PPP2R4*, as well as of *AURKA* and *PPP2CA*, was even more strongly associated with poor survival (Figure 4B). These data suggest that, as in the TH-MYCN mouse model, AURKA plays a role in neuroblastomagenesis despite concomitant transcriptional upregulation of the tumor suppressor *PPP2R4*.

**Figure 4:**
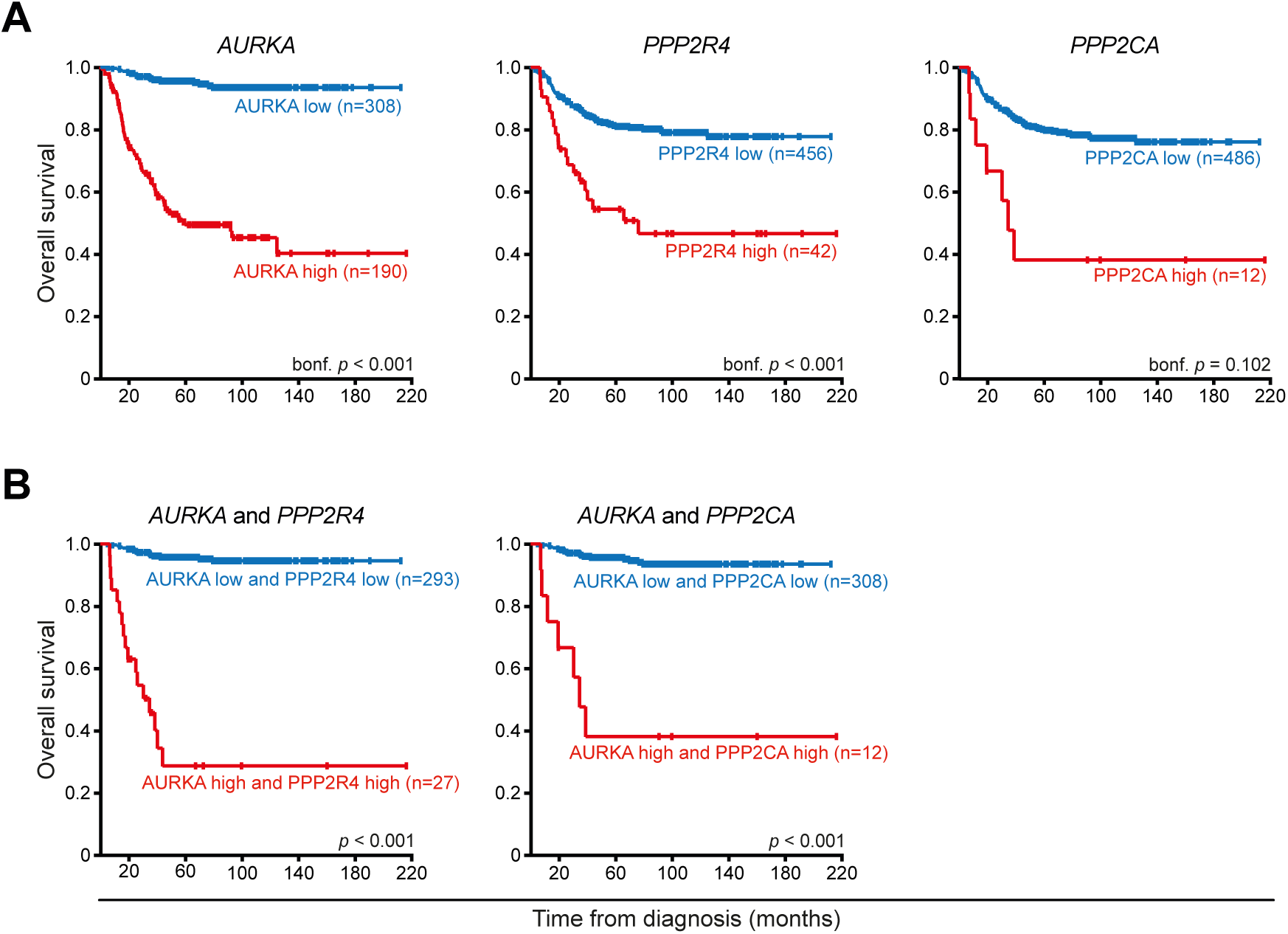
Enhanced expression of *AURKA* and *PPP2R4* is associated with poor prognosis in patient NB. Kaplan-Meier analysis of overall survival of 498 clinically annotated NB patients (SECQ-GSE62564) depending on transcript levels of *AURKA*, *PPP2R4* and *PPP2CA* **(A)** or of the combination of AURKA and PPP2R4 or AURKA and PPP2CA **(B)** are shown. Statistical analysis of Kaplan-Meier curves was performed using the log-rank test with Bonferroni correction. The cut-off was determined by the scanning method.

### Loss of *PPP2R4* decreases PPP2CA and increases AURKA protein in KELLY NB cells

Findings from the TH-MYCN neuroblastoma model and human NB samples suggested a role for PPP2R4 in regulating the AURKA network. To substantiate this, Bayesian analysis of microarray data from TH-MYCN tumors was performed using 85 genes important in AURKA signaling, identifying PPP2R4 as a central node in the network (Figure 5A). To experimentally assess the effect of PPP2R4 on AURKA, *PPP2R4* was knocked out in KELLY cells, and expression of methylated PPP2CA (mePPP2CA) and AURKA protein was determined. PPP2CA is the catalytic subunit of PP2A and its methylation promotes its function. Homozygous deletion of *PPP2R4*, confirmed by Sanger sequencing and Western blot, consistently decreased mePPP2CA expression and increased AURKA protein levels in all clones examined (Figure 5B).

**Figure 5:**
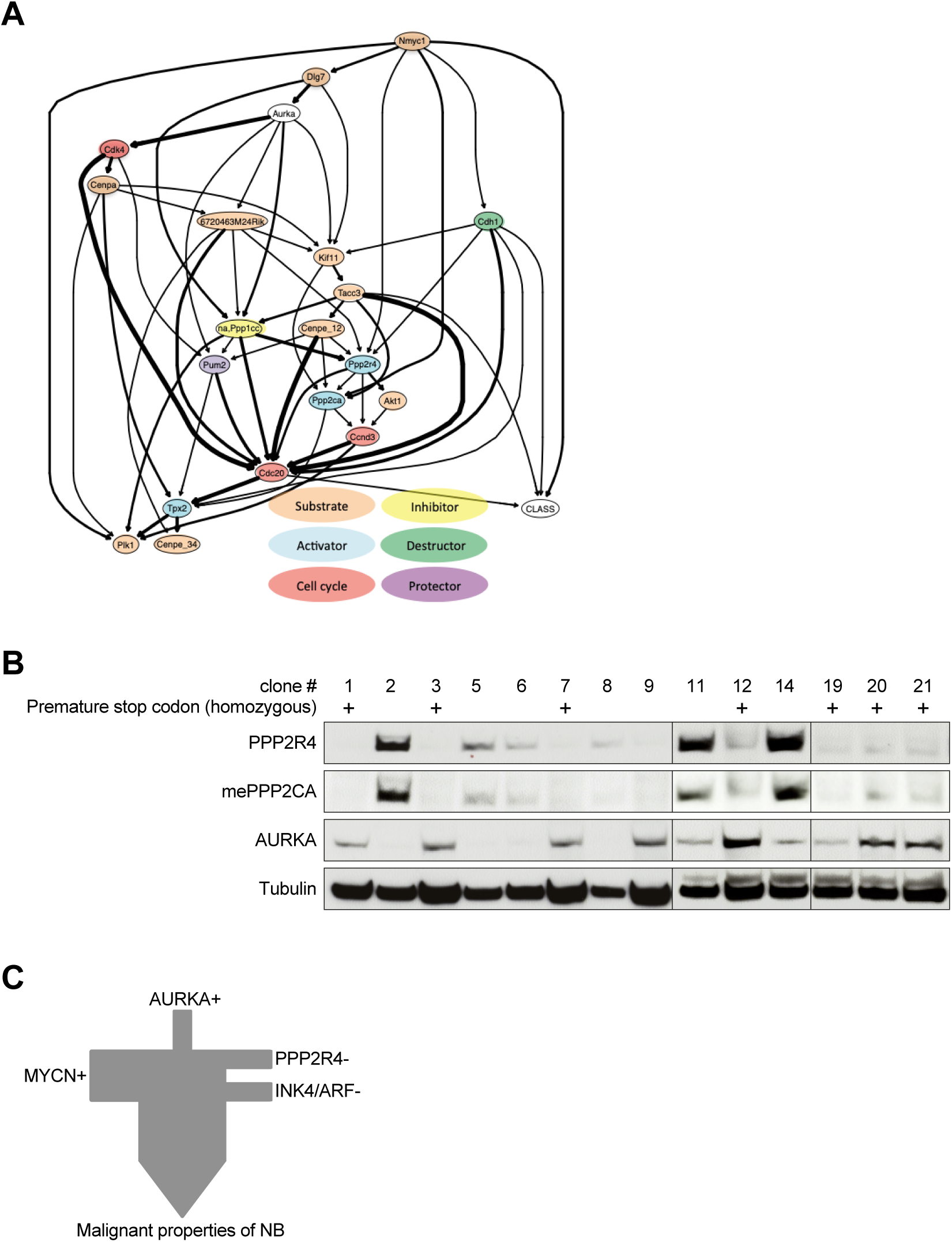
Loss of PPP2R4 increases AURKA protein in NB cells. **A) Bayesian analysis of mRNA expression in ganglia and NB of TH-MYCN mice suggests a role of PPP2R4 in dysregulation of AURKA in neuroblastomagenesis.** Microarray mRNA data previously described in (32) were used. 85 genes implicated in the AURKA network were subjected to Bayesian analysis. The direction (arrows) and probability (edge thickness) of influence of the top 20 genes according to the differences of mean expression in NB vs. ganglia are shown. Genes are color-coded according to function in relation to AURKA. **B) Loss of PPP2R4 depletes mePPP2CA protein and increases AURKA protein in KELLY cells.** KELLY cells were transiently transfected with a CRISPR/Cas9 construct targeting *PPP2R4*, sorted and seeded as single cells. Single-cell clones were expanded and classified as *PPP2R4* wild-type or homozygous knockout. Shown are Western blots of PPP2R4, methylated PPP2CA (mePPP2CA), AURKA and Tubulin as loading control. **C) Schematic cartoon depicting the relationship between AURKA, PPP2R4, MYCN and INK4A/ARF in promoting malignant properties of NB.**

In summary, PPP2R4 depletion increases AURKA and decreases PPP2CA protein in NB cells.

## Discussion

This study elucidates roles of AURKA in the initiation, progression, and maintenance of NB. We show that AURKA promotes transformation of MEFs only when MYCN is overexpressed and INK4/ARF is absent, promotes malignancy of established NB cells, and is negatively regulated by the PP2A pathway (summarized in Fig. 8C).

We first demonstrate that *AURKA* and *MYCN* mRNA expression declines physiologically in the murine adrenal gland after birth, possibly reflecting a reduction in SAPs. This downregulation supports a model where sustained or ectopic expression of AURKA and MYCN expands transformation-prone SAPs, initiating NB. However, whole adrenal glands contain many non-sympathetic cell types that may confound expression analyses.

AURKA, even in combination with MYCN, did not transform MEFs unless INK4A and ARF were deleted, indicating that G1 checkpoint disruption is necessary for AURKA-driven transformation. This aligns with studies in other cell types. AURKA transforms immortal Rat1 fibroblasts (36), which lack p53-dependent p21 (37), but not primary MEFs (10, 34). In NIH 3T3 cells, transformation occurs only under serum starvation (36, 38). These findings suggest AURKA requires a cooperating event, particularly p53 pathway impairment. Accordingly, AURKA-induced aneuploidy is worsened when p53 is impaired (39). Along this line, in CAG-loxP-CAT-loxP-AURKA mice, conditional overexpression in mammary glands does not cause tumors due to p53-mediated arrest and apoptosis (40). p53 loss in these mice prevents apoptosis but induces only benign tumors through senescence (41). In contrast, some studies report transformation by AURKA in mammary epithelium (42, 43). These differences may reflect varying promoter systems, expression duration, and cellular or tissue context.

To assess AURKA’s transforming ability in SAPs in vivo, we generated a transgenic mouse line expressing hAURKA in the peripheral sympathetic nervous system. *hAURKA* mRNA was detected in the adrenal medulla, but the protein was undetectable by immunohistochemistry. This was not due to technical failure, as AURKA was readily observed in human NB tissue, suggesting rapid degradation in normal SAPs, as observed in other cell types (44–46). In transgenic mice, AURKA is known to be degraded during the G1–S transition via the ubiquitin–proteasome system, explaining its absence despite transcription (47). These findings suggest AURKA protein must exceed a threshold to exert oncogenic effects and that normal SAPs have intrinsic mechanisms limiting its accumulation. The lack of transformation aligns with some studies in breast tissue (40, 41) while contradicting other (42, 43). The conflicting results may reflect differences in expression systems and cellular context.

To assess relevance in vivo, we analyzed gene expression in the TH-MYCN mouse model. *AURKA* mRNA expression rose early in NB development and increased during progression, paralleling *MYCN*. *CDKN2A* and *PPP2R4* mRNA expression also rose. While paradoxical, this may reflect a compensatory tumor-suppressive response. However, these mechanisms appear insufficient to halt NB, possibly due to post-transcriptional silencing or protein inactivation. These conclusions are limited by reliance on mRNA data, which do not reflect protein activity. Moreover, *CDKN2A* expression includes both INK4A and ARF, precluding isoform-specific analysis.

AURKA not only played a role in murine neuroblastomagenesis but also enhanced the malignancy of established human NB cells lacking *INK4A/ARF*. Thus, in SH-EP-MYCN-ER human NB cells, which are *CDKN2A*-null, moderate AURKA overexpression significantly increased MYCN-driven anchorage-independent growth, Mechanistically, AURKA binding has been shown to stabilize MYCN by preventing FBXW7-mediated degradation (10).

In patient NB datasets, high *AURKA* and *PPP2R4* mRNA expression correlated with poor prognosis, consistent with the TH-MYCN mouse findings and subject to similar caveats.

Given PP2A’s tumor-suppressive function in NB, we tested whether it modulates AURKA. Bayesian analysis of TH-MYCN mRNA data implicated PPP2R4 in dysregulating AURKA. Supporting this, *PPP2R4* knockout in KELLY cells reduced PPP2CA protein (the catalytic subunit of PP2A) and increased AURKA protein. This suggests PP2A, via PPP2R4, suppresses AURKA by promoting its dephosphorylation and degradation. Three points are notable: First, PPP2R4-deficient clones proliferated, indicating PPP2R4 is dispensable for KELLY cell survival, though this requires validation in other NB lines. Second, AURKA protein levels were lower in PPP2R4-expressing KELLY cells, likely due to PP2A stabilization (16) promoting AURKA degradation. This warrants confirmation in NB xenografts. Third, the apparent contradiction between increased *PPP2R4* mRNA in TH-MYCN and patient NB (associated with poor prognosis) versus its loss elevating AURKA protein is resolved by distinguishing mRNA from protein regulation.

In summary, AURKA promotes NB emergence and progression but requires impaired checkpoint function and oncogenic help to transform cells. MYCN and AURKA cooperate to sustain oncogenic signaling, while loss of INK4A/ARF or PP2A permits their accumulation and synergy. Reactivating PP2A or restoring G1 checkpoint control may offer therapeutic opportunities, as others and we have shown (17–19). AURKA protein declines when PPP2R4 and PPP2CA are expressed (this paper), and PP2A-mediated Ser51 dephosphorylation triggers AURKA degradation (21). These findings suggest AURKA is a downstream target of PP2A-activating drugs. Clinically, the AURKA inhibitor alisertib has shown promise in a phase 2 NB trial (48), and new molecules for targeted AURKA degradation are under development (49).

In conclusion, AURKA acts as an oncogenic driver in NB restrained by tumor suppressors, with PP2A serving as a key modulator of its function and expression.

## Acknowledgements

We thank Dominik Stadel for providing data. We are very grateful to Glenn M. Marshall, Daniel R. Carter, Bellamy B. Cheung, Katleen De Preter and Aneleen Beckers for sharing and interpreting data from the TH-MYCN progression model, and to Glenn M. Marshall for valuable general discussions. The excellent technical assistance of Helgard Knauss is greatly appreciated.

This work was funded in part by grants of the Deutsche Forschungsgemeinschaft (DFG) to C.B. (B1-GRK2254 and BE 2416/12-1). G.L.S. received support of the International Graduate School in Molecular Medicine, Ulm.

## Author contributions

C.B. conceived and designed the experiments. G.L.S., M.D. and H.K. performed the experiments, analyzed the data and interpreted the results. All authors discussed the results. G.L.S. and C.B. wrote the manuscript. All authors read and approved the final manuscript.

